# Evaluation of the speed, accuracy and precision of the QuickMIC rapid antibiotic susceptibility testing assay in a clinical setting

**DOI:** 10.1101/2021.08.11.455925

**Authors:** Christer Malmberg, Jessie Torpner, Jenny Fernberg, Håkan Öhrn, Cecilia Johansson, Thomas Tängdén, Johan Kreuger

## Abstract

The rapidly changing landscape of antimicrobial resistance poses a challenge for empirical antibiotic therapy in severely ill patients and highlights the need for fast antibiotic susceptibility diagnostics to guide treatment. Traditional methods for antibiotic susceptibility testing (AST) of bacteria such as broth microdilution (BMD) or the disc diffusion method (DDM) are comparatively slow and show high variability. Rapid AST methods under development often trade speed for resolution, sometimes only measuring responses at a single antibiotic concentration. QuickMIC is a recently developed lab-on-a-chip system for rapid AST. Here we evaluate the performance of the QuickMIC method with regard to speed, precision and accuracy in comparison to traditional diagnostic methods. 151 blood cultures of clinical Gram-negative isolates with a high frequency of drug resistance were tested with the QuickMIC system and compared with BMD for 12 antibiotics. To investigate sample turnaround time and functionality in a clinical setting, another 41 clinical blood culture samples were acquired from the Uppsala University Hospital and analyzed on site in the clinical laboratory with the QuickMIC system, and compared with DDM for 8 antibiotics routinely used in the clinical laboratory. The overall essential agreement between MIC values obtained by QuickMIC and BMD was 83.2%, with an average time to result of 3 h 2 min (SD: 24.8 min) for the QuickMIC method. For the clinical dataset, the categorical agreement was 94.9%, and the total turnaround time as compared to routine diagnostics reduced by 40% (33 h vs. 55 h). Interexperiment variability was low (average SD: 44.6% from target MIC) compared to the acceptable standard of ±1 log_2_ unit (*i*.*e*. -50% to +100% deviation from target MIC) in BMD. We conclude that the QuickMIC method can provide rapid and accurate AST, and may be especially valuable in settings with high resistance rates and for antibiotics where wildtype and resistant MIC distributions are close or overlapping.

## 1 Introduction

Sepsis has recently been recognized as the most common cause of death globally, with an estimated 48.9 million sepsis cases each year, resulting in approximately 11 million deaths^1^. Rapid administration of adequate antibiotics is key to efficient treatment of severe bacterial infections that otherwise can lead to life-threatening conditions such as sepsis or septic shock^2,3^. According to international guidelines for management of sepsis and septic shock, administration of antibiotics should be initiated within 1 hour from recognition of sepsis ^4^, and a number of studies show that early administration of effective therapy results in increased clinical as well as financial benefits^3,5– 7^. However, the escalating prevalence of multidrug-resistant bacteria worldwide reduces the probability of empirical antibiotic therapy being microbiologically active^4^. Thus, rapid diagnostics of antibiotic susceptibility is becoming increasingly important to avoid treatment failure. In recent years there has been an increasing focus on achieving faster clinical microbiology diagnostics such as rapid bacterial identification (rID) and rapid antibiotic susceptibility testing (rAST). Routinely used AST methods like broth microdilution (BMD) and the disc diffusion method (DDM) can be made faster, both by improved diagnostic logistics but also as a result of updated and improved protocols. Furthermore, EUCAST recently presented an improved protocol including new breakpoints for rapid read-out of disc diffusion (to be used within 4, 6 and 8 hours of plate inoculation), which indicates the possible speed gains from updated protocols^8^. However, the actual benefits of the more rapid, traditional methods developed so far has been questioned^9,10^, due to problems with uneven performance and analysis times still being too long.

Notably, low- and-middle income countries are disproportionally burdened by sepsis mortality, much due to the lack of effective healthcare systems, including diagnostic services. Access to effective diagnostics has been argued to be as fundamental as access to antibiotics^11–13^, but remains problematic in resource-poor settings due to the lack of appropriate infrastructure needed to effectively implement diagnostic guided therapy. Hopefully addressing this problem are new, affordable, automated and networked diagnostic systems capable of being operated in an outpatient setting, and not necessarily by laboratory professionals. This is a direction which has been investigated in several new diagnostic systems recently reaching the market, especially with a focus on molecular diagnostics for rID and rapid resistance screening. Examples include the GenMark ePlex® system, Curetis Unyvero™ and the BioMerieux FilmArray® systems^14^. However, to our knowledge there are currently no approved AST diagnostic systems which fit these criteria.

Based on these considerations we have previously developed a new microfluidic rAST method^15,16^ where bacterial responses are monitored in precisely controlled antibiotic gradients. We have demonstrated the use of this system for rapid AST directly from positive blood cultures, for a single channel prototype system as well as a proof-of-concept multiplex system. In this study we investigated the performance of a refined system based on our earlier design, called QuickMIC (Figure 1). The ultimate goal is to develop QuickMIC into a rapid, automated and affordable diagnostic solution for AST, with potential to expand access to rapid diagnostics, and this new rAST diagnostic system is currently available for research use only (RUO) applications. The aim of this study was to evaluate the performance of the current RUO system with respect to accuracy, precision, time-to-result; as well as the feasibility of using the QuickMIC system in a clinical setting. This study aimed to generate indicators for use in fine-tuning the system performance, and can be seen as a pilot study for further evaluation purposes. Briefly, we show that the QuickMIC system can reduce sample turnaround times in a clinical setting with at least 40% compared to disc diffusion, with high categorical agreement (94.9%) to standard testing.

**Figure 1:**
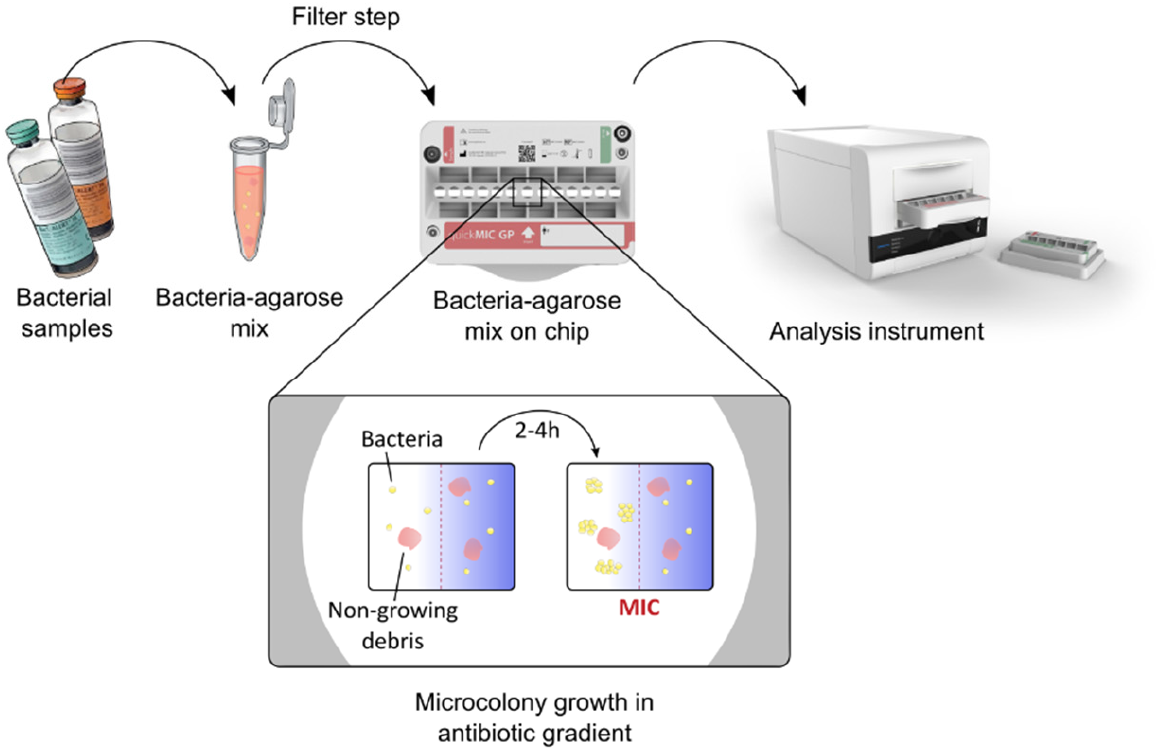
Overview of the QuickMIC system and the sample preparation process. In short, blood culture samples are mixed with a stabilizing agarose matrix, filtered to remove blood cells and injected into the QuickMIC cassette. The cassette is then inserted into an analysis instrument, where bacterial growth in an array of growth chambers containing polymerized agarose gels is monitored in the presence of a predetermined set of linear antibiotic gradients. Microcolony growth is evaluated using time-lapse imaging for up to 4 hours, and stable antibiotic inhibition zones are typically detectable after 2-4 h and used to automatically calculate MIC values for the different antibiotics.

## 2 Material and methods

### 2.1 Handling of bacterial isolates for reference testing

A total of 151 bacterial isolates (species distribution detailed in Table 1) were acquired from several sources, with varying levels of preexisting knowledge of the antibiotic susceptibility profile for each strain. The sources were the EUCAST development laboratory (EDL, Växjö), the Antibiotic Research Unit, Uppsala University and Uppsala University Hospital. The bacterial strains were streaked on arrival, grown overnight and thereafter kept frozen in -70°C for the remainder of the study. All strains were cultivated using Müller-Hinton (MH-II, BBL, Becton Dickinson) agar and cation-adjusted MH-II broth. For QuickMIC and BMD testing, suspensions of bacteria were prepared by dissolving two to four colonies in MH-II, after which the suspension was adjusted to 0.5 McFarland and either used for BMD or diluted and inoculated into a BacT/Alert FA Plus bottle (bioMerieux, Marcy l’Etoile, France) together with 10 mL horse blood, to an initial concentration of either 25 cfu/mL or 2.5*10^5^ cfu/mL. The blood culture bottles were incubated in a BacT/Alert Virtuo system (bioMerieux) until a positive signal was received, after which the bottle was removed for QuickMIC testing. The timepoint when the bottle was removed from the BacT/Alert system for testing was recorded.

**Table 1:** Included species and strain numbers in the reference panel.

| Species | Number of different strains per species | % of all evaluated species |
| --- | --- | --- |
| *A. baumannii* | 20 | 13.2 |
| *C. koseri* | 8 | 5.3 |
| *E. aerogenes* | 8 | 5.3 |
| *E. cloacae* | 14 | 9.3 |
| *E. coli* | 30 | 19.9 |
| *K. oxytoca* | 5 | 3.3 |
| *K. pneumoniae* | 30 | 19.9 |
| *P. aeruginosa* | 17 | 11.3 |
| *P. mirabilis* | 9 | 6.0 |
| *S. marcescens* | 10 | 6.6 |
| **Total:** | 151 | 100 |

On-scale quality-control (QC) strains were used during the study to repeatedly track the QuickMIC method performance, as per the instructions for use available from the manufacturer (Gradientech AB, Uppsala, Sweden). Results from these continuously performed runs were pooled and intra-laboratory variability calculated for quantification of method repeatability.

### 2.2 Broth microdilution AST

Broth microdilution was performed according to ISO 20776-1:2019 (International Organization for Standardization [ISO]). The antibiotics and concentrations used are described in Table 2. Briefly, the microplates (Greiner Bio-one, flat bottom PS, product number: 655101) were loaded with antibiotics and inoculated with a bacterial suspension at 0.5 McFarland, yielding a final concentration of ∼5*10^5^ CFU/mL; after which they were incubated overnight at 37°C. The MIC was read after 16 h as the lowest drug concentration at which no turbidity could be detected visually. All strains were tested at least twice on separate days, in case of a differing result the strain was retested a third time and the MIC reported as the mode of the three read-outs.

**Table 2:** Antibiotic concentrations used for BMD and QuickMIC AST testing.

| Antibiotic (#) | Supplier | Article number | Batch number | Concentration range mg/L (QuickMIC) | Concentration range mg/L (BMD) |
| --- | --- | --- | --- | --- | --- |
| Amikacin (AMI) | Sigma | PHR1860 | LRAB1258 | 1 – 20 | 0.5-32 |
| Cefepime (CEP) | Sigma | PHR1763 | LRAB8503 | 0.5-10 | 0.25-16 |
| Ciprofloxacin (CIP) | Sigma | PHR1044 | LRAB3671 | 0.125-2.5 | 0.125-4 |
| Colistin (COL) | Sigma | C4461 | 089M4881V | 0.25-5 | 0.125-8 |
| Cefotaxime (CTA) | Sigma | Y0000420 | 4.0 | 0.25-5 | 0.0156-8 |
| Ceftazidime / (CTZ) | Sigma | C0690500 | 3 | 0.5-10 | 0.25-16 |
| Avibactam (CTV) | AmBeed | A169351 | A169351-002 | 4* | 4* |
| Gentamicin sulfate (GEN) | Sigma | G4918 | 028M4827V | 0.5-10 | 0.25-16 |
| Meropenem (MER) | Sigma | PHR1772 | LRAB7853 | 0.5-10 | 0.25-16 |
| Piperacillin / | Sigma | PHR1805 | LRAB7665 | 2-40 / | 0.5-64 / |
| Tazobactam (PIT) | Sigma | 68300 | BCCD0509 | 4* | 4* |
| Tigecycline (TIG) | Sigma | Y0001961 | 1.0 | 0.06-1.25 | 0.0156-2 |
| Tobramycin (TOB) | Sigma | T1783 | SLBX8043 | 0.5-10 | 0.125-16 |

### 2.3 QuickMIC rapid AST

For QuickMIC testing, samples from positive blood culture bottles were prepared according to the instructions for use available from the manufacturer (Gradientech AB, Uppsala, Sweden). In short, 2-4 mL of blood culture was aspirated using an S-Monovette safety needle (Sarstedt, Nümbrecht, Germany), after which a 10 μL sample loop was used to transfer material to the QuickMIC preparation kit (art. nr: 46-001-10). The preparation kit performs a fixed 1:200 dilution and filtering step to remove blood cells. An aliquot of prepared sample was kept for inoculate control by plating. The sample was then loaded in the QuickMIC test cassette (GN panel, art. nr: 43-001-10) sample port, after which 20 mL of MH-II was injected in the cassette media port. The cassette was inserted into the instrument and the test was started through the control software (QM Analyst v0.93). The QuickMIC RUO (v0.93) system was used for the entirety of this study, in this case consisting of 6 instrument modules connected to a control PC. In the current study, the QuickMIC GN cassettes were hand loaded with antibiotics according to the concentrations in table 2, by the following procedure: Each antibiotic well was loaded with 49 μl stock concentration of 10x the concentration indicated in Table 2, which was then diluted to final concentration during the MH-II loading step. Each of the 151 strains were tested once using the QuickMIC system.

### 2.4 AST of clinical blood cultures

A total of 48 clinical samples were collected as per the following protocol. After indication of growth in the BactAlert Virtuo system and indication of Gram-negative bacteria, 2 mL samples from the blood culture bottles (FA Plus or FN Plus) were provided by the Department of Clinical Microbiology at Uppsala University Hospital. The samples were labelled with a sample code for anonymization, and an aliquot of the sample was streaked on MH-II agar for determination of inoculate concentration, as well as for isolation of the strain. The remaining sample was prepared identically as per the previous section for QuickMIC rAST analysis. Single colonies from the streaked plate were used to create a frozen stock for BMD testing, which was then performed as described above. Afterwards, for all tested strains the bacterial identity (MALDI Biotyper, Bruker Daltoniks), susceptibility category by disc diffusion and data on test dates and times to result were provided by the clinical microbiological laboratory. Only samples where non-facultative Gram-negative pathogens where identified as single species were approved for further data analysis (*n* = 41). After test completion, or at longest 6 months after sample collection, the original blood samples were destroyed and the dataset was de-identified by destruction of the code key. The study protocol for the clinically derived samples was approved by the Swedish Ethical Review Authority (Dnr 2020-03060).

### 2.5 Data analysis

The MIC values generated by the QuickMIC system (QM Analyst v0.93) were compared with BMD values as previously described^15^. In short, linear-scale MIC values were right-censored to nearest log2 dilution step, and essential agreement counted as within 1 log2 step of the corresponding BMD result. When the reference method showed results below or above the limit of quantification (LOQ) for the tested method, these were counted as in agreement. For comparison of categorical agreement, susceptible (S) instead of resistant (R) was counted as a very major error (VME), R instead of S as a major error (ME) and S or R instead of increased exposure (I) or vice versa as a minor error (MiE). The categorization was performed by applying EUCAST clinical breakpoints (version 10.0, 2020, available at www.eucast.org). Data on times to result and turnaround times for each analysis performed were compared using the freely available Jamovi distribution of R (v1.68). For statistical analysis of parameters influencing the result quality, logistic and linear regression was performed using Jamovi.

## 3 Results

### 3.1 QuickMIC performance from spiked blood samples

After acquiring a reference strain collection of 151 nonfacultative Gram-negative strains of clinical origin (Table 1) we performed rapid AST testing using the QuickMIC system to assess the performance of the novel assay in comparison with BMD. Each strain was tested with 12 antibiotics, as detailed in Table 2. A positive control chamber not exposed to antibiotics is included in the QuickMIC cassette, and only cassettes with positive control growth and channels where mean fluidic flow was within ±50% of nominal flow (1 μl/min) at the end of the experiment were accepted as valid runs (*n* = 1671, 92% of dataset). MIC values were obtained from the QuickMIC system data analysis algorithm (v1), and all MIC values from valid runs were compared to reference MICs from BMD (Table 3). During the study, three on-scale QC strains (*E. coli* NCTC 13846, *K. pneumoniae* ATCC 700603 and *P. aeruginosa* ATCC 27853 were regularly run with the system to track performance. These replicate runs were performed on separate days (*E. coli n* = 21, *K. pneumoniae n* = 19, *P. aeruginosa n* = 22), and were pooled to measure repeatability of the QuickMIC system.

**Table 3a.** Overall essential agreement and categorical agreement between QuickMIC and BMD for the reference strains, by tested antibiotic.

| *n* = | Antibiotic |  |  |  |  |  |  |  |  |  |  |  |  |
| --- | --- | --- | --- | --- | --- | --- | --- | --- | --- | --- | --- | --- | --- |
|  | AMI<br>(96) | CEP<br>(139) | CIP<br>(144) | COL<br>(147) | CTA<br>(136) | CTV<br>(131) | CTZ<br>(139) | GEN<br>(145) | MER<br>(151) | PIT<br>(136) | TIG<br>(138) | TOB<br>(133) | Total<br>(1635) |
| EA (%) | 70.8 | 84.2 | 91.0 | 71.4 | 86.0 | 89.3 | 75.5 | 91.7 | 84.1 | 81.6 | 87.7 | 82.0 | 83.2 |
| CA (%) | 94.9 | 81.7 | 83.0 | 92.6 | 86.9 | 99.1 | 72.5 | 96.9 | 84.9 | 78.5 | 48.5 | 89.8 | 86.3 |

**Table 3b.**
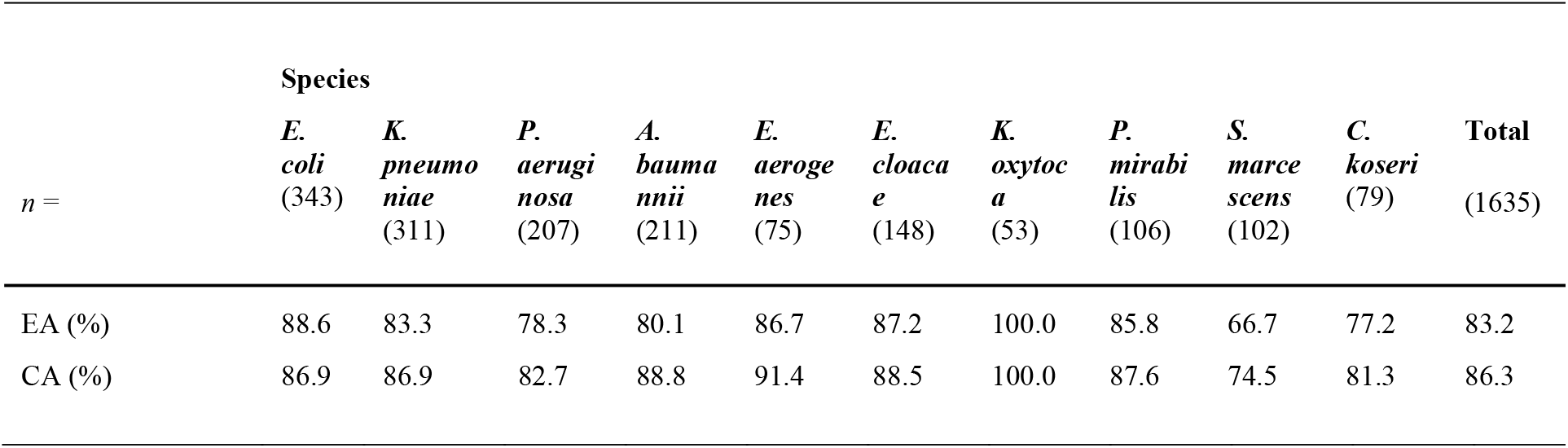
Overall essential agreement and categorical agreement between QuickMIC and BMD for the reference strains, by tested species.

Of 1671 valid QM results, only 1635 had a reference BMD value for comparison. For the antibiotics tested, the essential agreement (EA) ranged from 70.8% to 91.7% (mean: 83.2%, *n* = 1635) depending on antibiotic (Table 3a) and from 66.7 to 100% depending on species tested (Table 3b). The overall categorical agreement (CA) for the dataset was 86.3%, ranging from 48.5 to 99.1% depending on antibiotic tested, and from 74.5 to 100% depending on species tested. Categorical agreement was only calculated for drug and bacteria combinations with breakpoints as established by EUCAST. The distribution of errors for each antibiotic for the 4 most common Gram-negative bacterial species isolated in blood stream infections can be seen in Figure 2.

**Figure 2.**
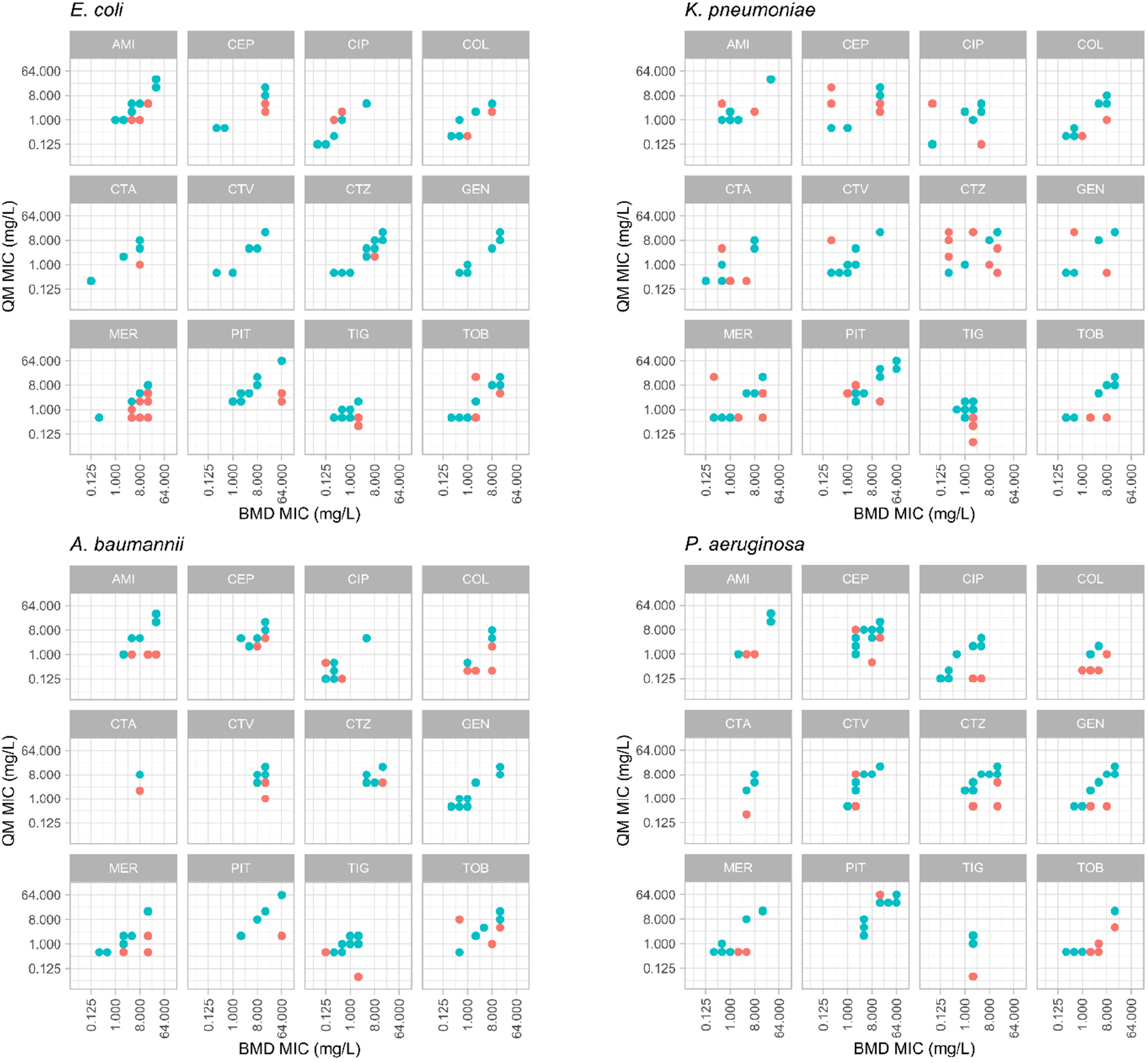
Distribution of MIC value results from reference BMD method (x-axes) and the QuickMIC (QM) method (y-axes), for the 4 most common bacterial species encountered in blood stream infections, in response to the 12 tested antibiotics. MIC-values obtained by the QuickMIC method that are within EA in comparison to BMD are shown in blue, MIC values not in agreement are shown in red. AMI: amikacin, CEP: cefepime, CIP: ciprofloxacin, COL: colistin, CTA: cefotaxime, CTV: ceftazidime/avibactam, CTZ: ceftazidime, GEN: gentamicin, MER: meropenem, PIT: piperacillin/tazobactam, TIG: tigecycline, TOB: tobramycin.

The QuickMIC system reports individual MIC values from the cassette as soon as they are stabilized, which is called after that the MIC read-out value has fluctuated less than 5% over 30 minutes. The average time to result for the reference dataset was 182 min (SD: ± 24.8 min), with a range from 150 to 230 min.

### 3.2 Method repeatability

In total, the QC strains were tested on 62 occasions during the 8-month study. The three QC strains were selected to together give an on-scale result for all antibiotics included in the panel (except meropenem, where no on-scale ATCC QC-strain could be found), and were used to track method performance as per the instructions for use for the QuickMIC system. This dataset also allows a quantitative determination of method repeatability. The linear-scale MIC results are shown in Figure 3, relative to the reference MIC of the strain and the acceptable ±1 log2-step variation. One strain per antibiotic was used for QC testing (Figure 3, black arrow). The method repeatability for each antibiotic, normalized to target MIC for the respective QC strain, is shown in table 4. The average repeatability (mean SD) was 44.6% of the target MIC value, with a range from 13.4 to 150.5%. The highest repeatability was displayed by ceftazidime and the lowest by gentamicin. The acceptable BMD variation corresponds to -50% to +100% variation from the target MIC value on a linear scale, meaning that all antibiotics except ceftazidime had acceptable variability in results.

**Table 4:**
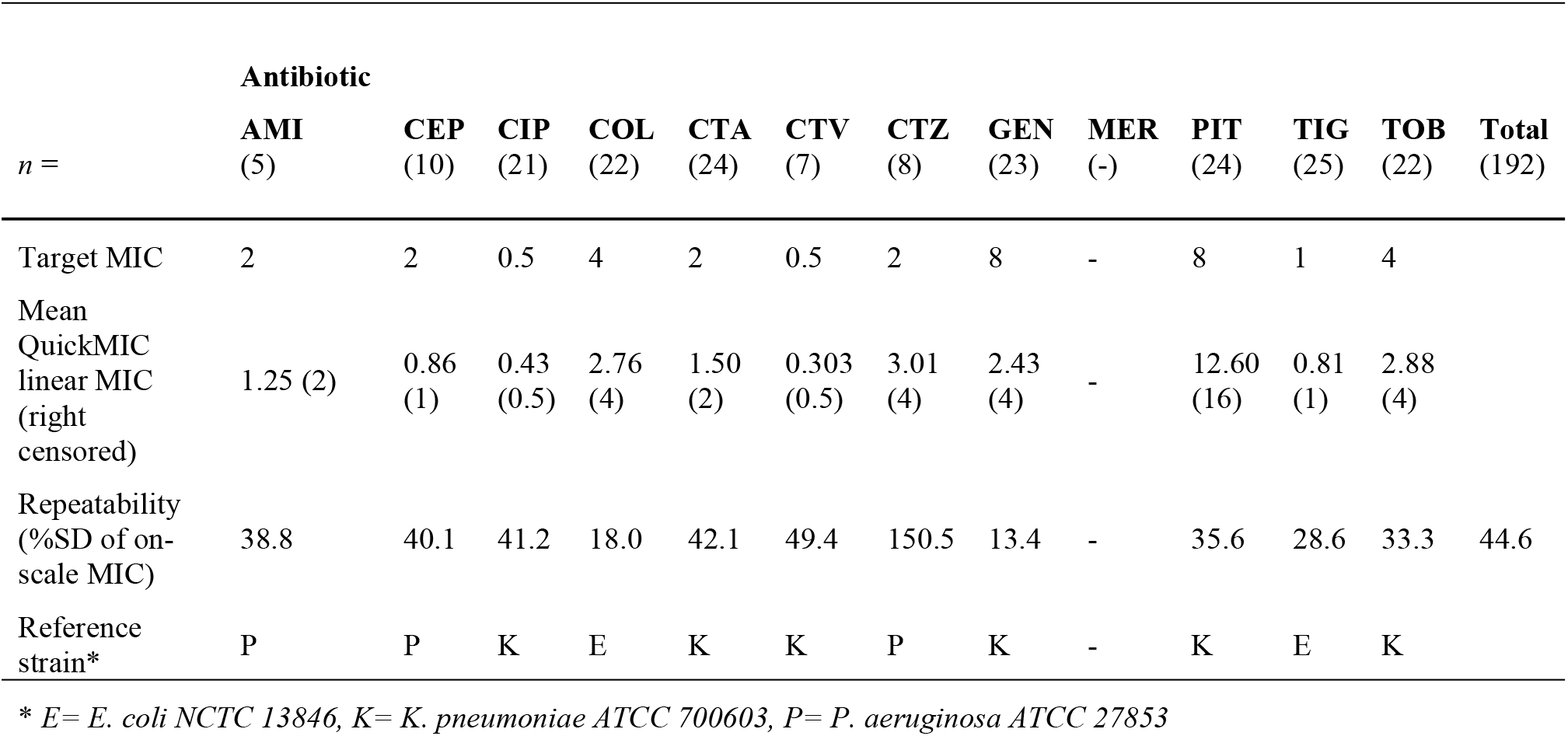
Repeatability per antibiotic for each panel reference strain. Only on-scale results were used for repeatability calculations.

**Figure 3:**
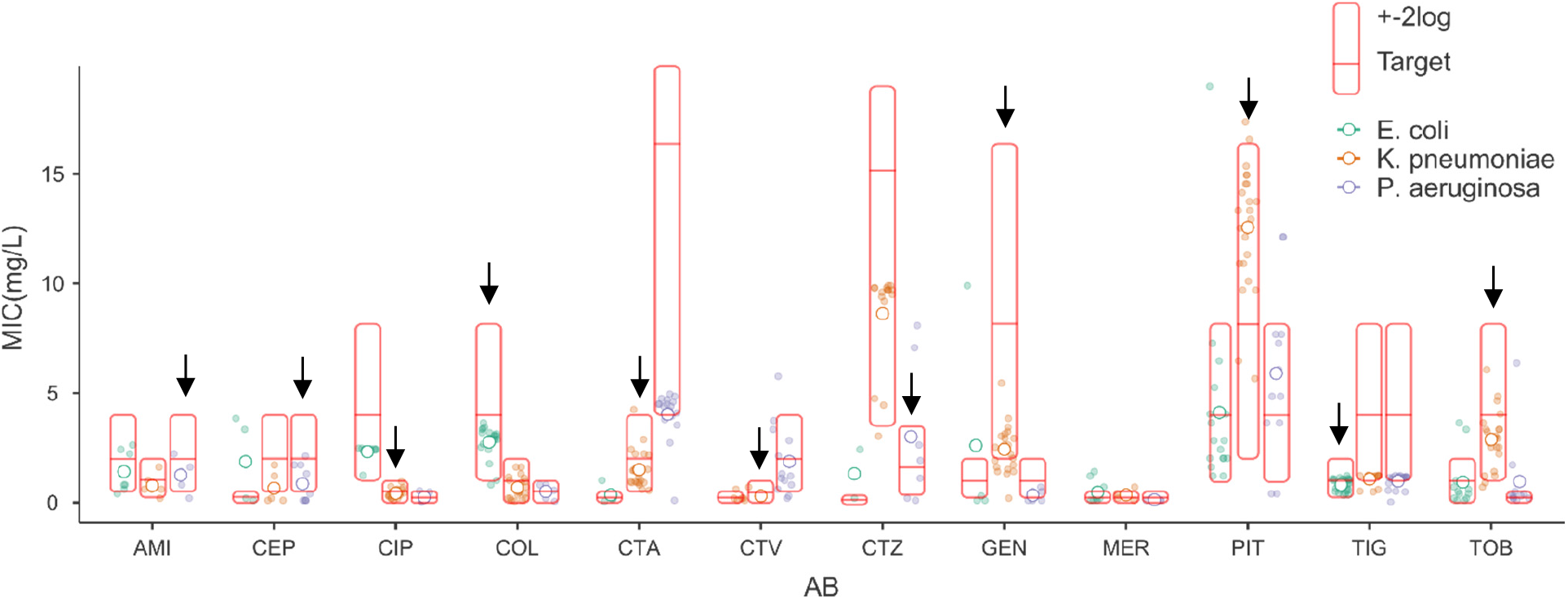
Repeatability per tested antibiotic for the three QC strains. The linear-scale MIC output (i.e. not right-censored for direct comparison with BMD) from the QuickMIC instrument is shown relative to the reference BMD MIC (red horizontal bar), with allowable variation from BMD reference (red box). Open circle: mean value for each antibiotic. Closed circles: raw data. Black arrow: on-scale QC strain for this antibiotic. Note that to compare accuracy with BMD, the QuickMIC linear-scale value should be right censored to the next log2 dilution.

### 3.3 Performance of the sample preparation process

The QuickMIC system is designed to be robust with regards to initial inoculate concentration and species, since in a clinical setting the actual bacterial concentration in the blood culture is unknown. In the reference dataset, 94.7% of blood cultures after sample processing yielded an inoculate concentration within the stated inoculate limits of the QuickMIC system (5*10^4^ cfu/mL to 5*10^8^ cfu/mL). As blood incubation systems are highly sensitive, at positivity the bacteria have not yet reached stationary phase and will continue to grow for several hours. If a bottle turns positive during off-shift hours in a clinical situation, it could be sampled hours later. To get a realistic range of start inoculates, the blood culture bottles were started at varying timepoints of the day. The time after positivity until the blood culture bottle was sampled ranged from 0 h – 10 h, and the inoculate concentration after sample preparation ranged from 8*10^3^ to 3.7*10^7^ cfu/mL. The correlation between time after positivity and starting inoculum was investigated by linear regression, and determined to be close to zero (slope 0.075 log(cfu/mL)/h, 95% confidence interval 0.06 – 0.08).

### 3.4 Parameters affecting the performance of QuickMIC

Rapid AST methods such as the QuickMIC assay must support variations in sample properties, since the start material is unprocessed blood culture material. Parameters that can be assumed to affect the analysis are total incubation time of the blood culture (i.e. incubation time plus waiting time until the positive signal is acted on), bacterial species, composition of the blood, and actual inoculate concentration. Also, the antibiotic and level of susceptibility can be important determinants, where some antibiotics and resistance mechanisms are more difficult to measure rapidly. To investigate some of these parameters, logistic regression was performed on the spiked blood sample dataset with overall EA as dependent variable, and species, antibiotic, inoculate concentration and inoculation time after alarm used as factors. The results of the regression are presented in Table 5. As expected, antibiotic, species and resistance category were significant factors influencing the performance of the method. Neither inoculate time after positivity (p = 0.951) nor inoculate concentration (p = 0.111) were significant factors however; the predicted impact of these parameters can be seen in Figure 5.

**Table 5.**
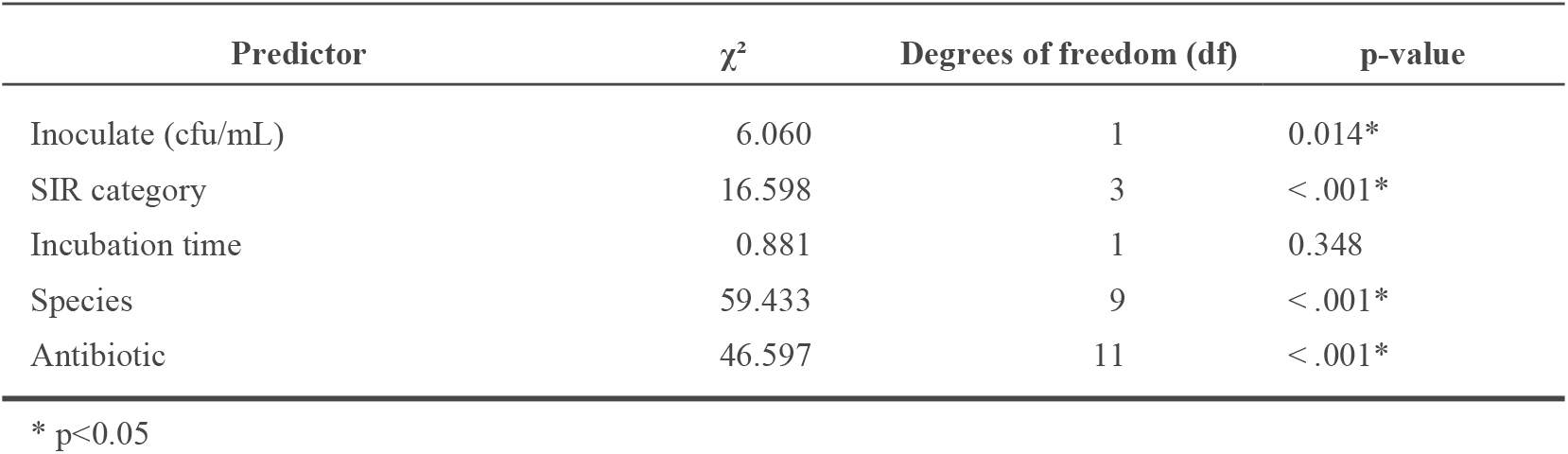
Logistic regression model of important parameters affecting EA

| Predictor | $\chi^2$ | Degrees of freedom (df) | p-value |
| --- | --- | --- | --- |
| Inoculate (cfu/mL) | 6.060 | 1 | 0.014* |
| SIR category | 16.598 | 3 | <.001* |
| Incubation time | 0.881 | 1 | 0.348 |
| Species | 59.433 | 9 | <.001* |
| Antibiotic | 46.597 | 11 | <.001* |

\* p<0.05

**Table 6.** Categorical agreement of QuickMIC and rapid disc diffusion in a clinical setting.

| Antibiotic<br>$n_{\text{=}}$ | AMI<br>(17) | CIP<br>(33) | CTA<br>(39) | CTZ<br>(39) | GEN<br>(33) | MER<br>(39) | PIT<br>(39) | TOB<br>(39) | Total<br>(893) |
| --- | --- | --- | --- | --- | --- | --- | --- | --- | --- |
| Categorical agreement (%) | 88.2 | 97.0 | 95.1 | 83.3 | 98.1 | 97.4 | 94.9 | 81.0 | 94.9 |

**Figure 4:**
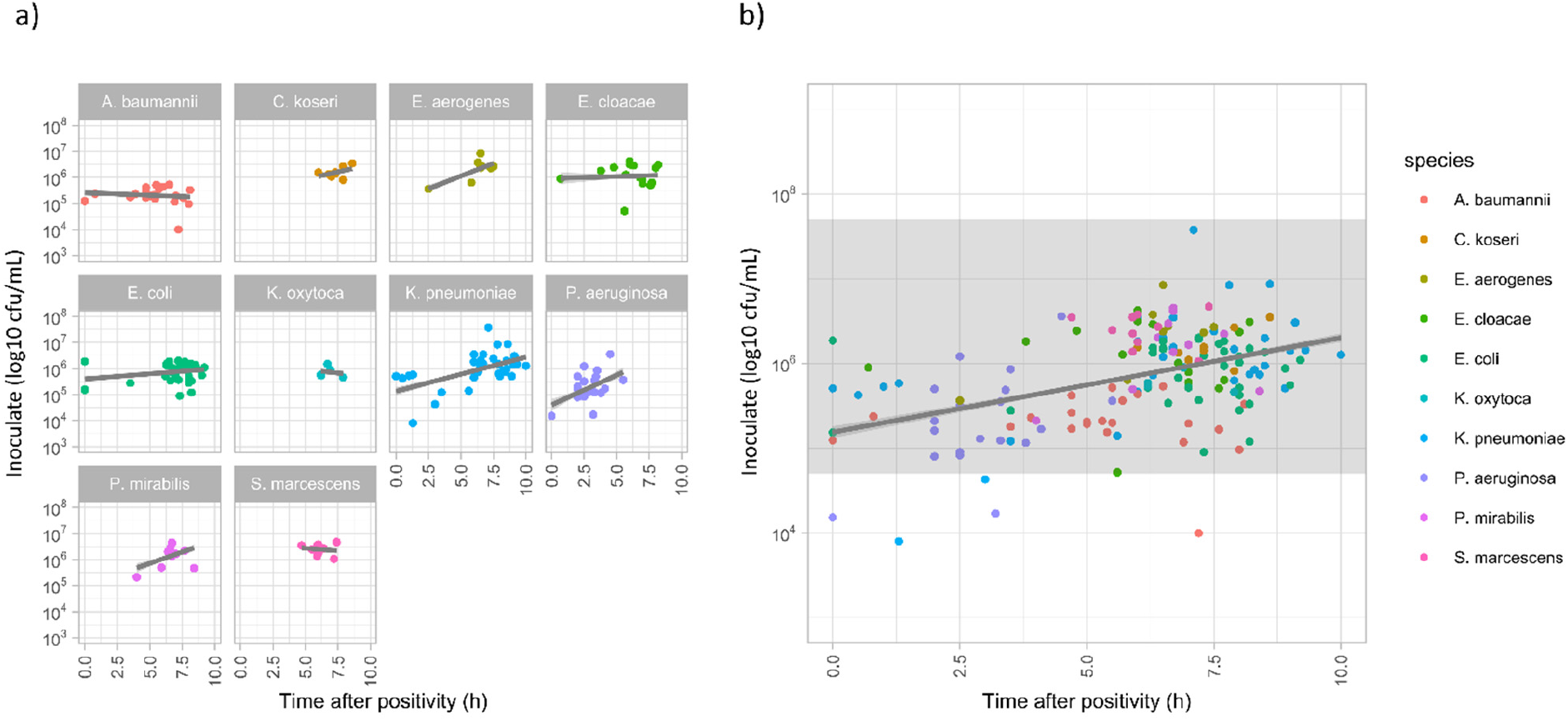
a) Bacterial inoculates (log10 cfu/mL) after sample preparation in comparison to the time (h) from positivity signal of the blood culture bottles to sampling. Grey line: linear regression line for each species. No specific correlation can be seen overall. b) All inoculates and supported range (grey field) for the QuickMIC system.

**Figure 5.**
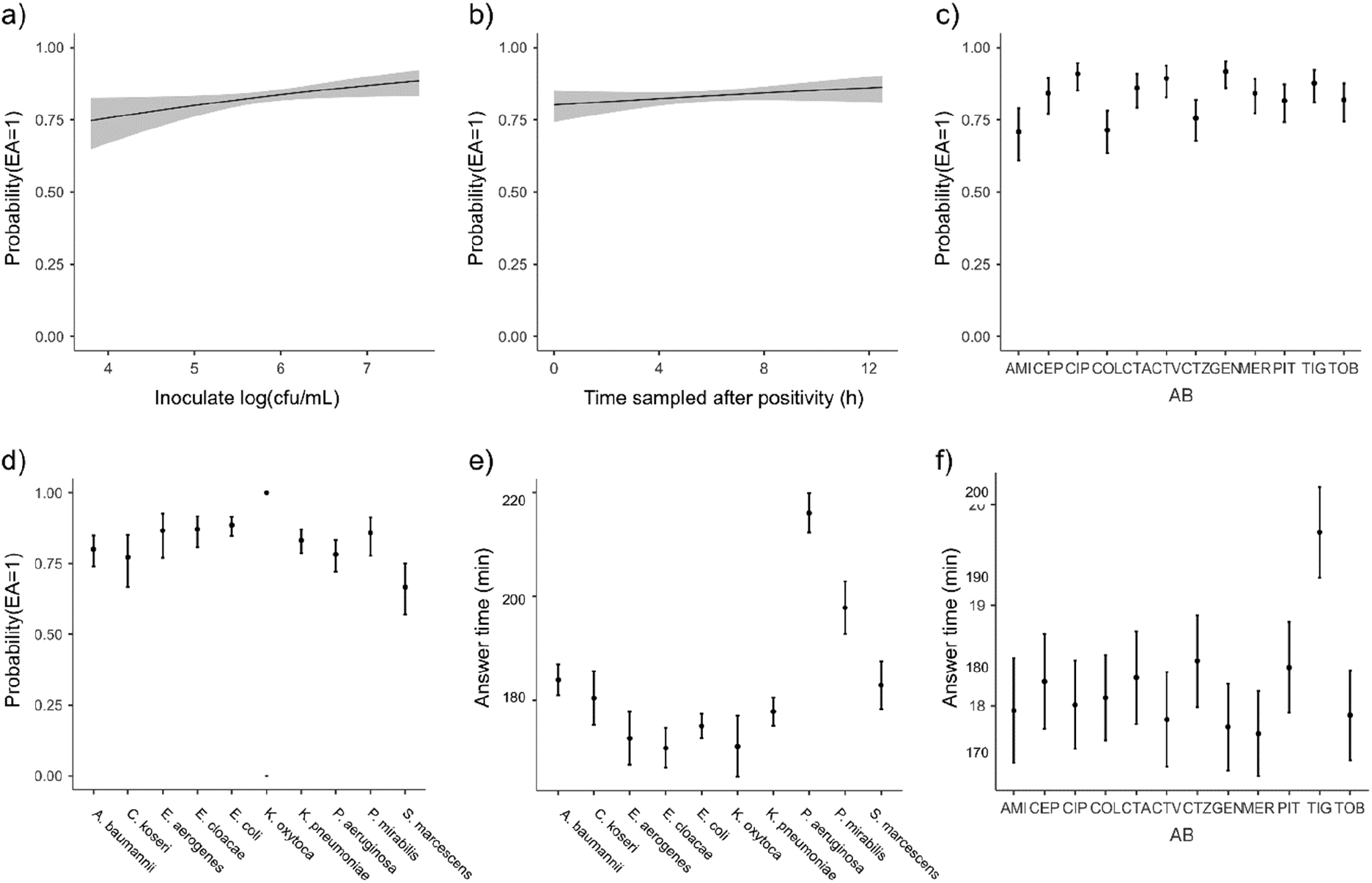
Performance affecting parameters. a-d) Logistic regression of inoculate concentration, time in incubator after positivity, species and antibiotic against EA. P(EA = 1): probability to achieve a result within EA as compared to BMD in the dataset from spiked blood cultures, as estimated by regression analysis. Grey fields, error bars: 95% confidence interval of prediction. e-f) Linear regression of species and antibiotic factors against answer time from the QuickMIC system.

### 3.5 Performance with clinical samples directly in a hospital microbiology lab setting

To test the capability of the method in a clinical setting, blood culture samples were collected from the Department of Clinical Microbiology, Uppsala University Hospital (*n* = 48). 41 samples (85%) were analyzable on the QuickMIC system (monomicrobial, nonfastidious Gram-negative bacteria within the supported concentration interval). The results showed that QuickMIC has 94.9% CA with DDM, as performed by the clinical laboratory. The mean time to result for each drug and bacteria combination was 178 min (SD: ±22.5 min). The samples were collected during early morning and run in the afternoon on the same day. To investigate the impact on total turnaround-time for the QuickMIC method as compared to the clinically used routine method, the time from patient sampling until test result availability (turnaround time, TAT) was recorded for QuickMIC and compared with turnaround-time values for DDM as per data from the clinical laboratory (Figure 6). In total, the mean turnaround time was reduced by 40%, and determined to be 33.4 h (SD: 13.0 h) for QuickMIC vs. 55.4 h (SD: 25.2 h) for DDM. The fastest QuickMIC total turn-around time was 21.3h, while the fastest DDM result was 36.1h. For the QuickMIC system the total turnaround-time was categorized into “transport”, “blood culture”, and “analysis” times. The transport time accounted for 48.7% (SD: 15.2%) of the turnaround time, whereas blood culture time (the time from insertion to positive alarm) was 41.3% (SD: 15.3%), and the time for QuickMIC analysis corresponded to 10.0% (SD: 3.0%) of the turnaround time.

**Figure 6:**
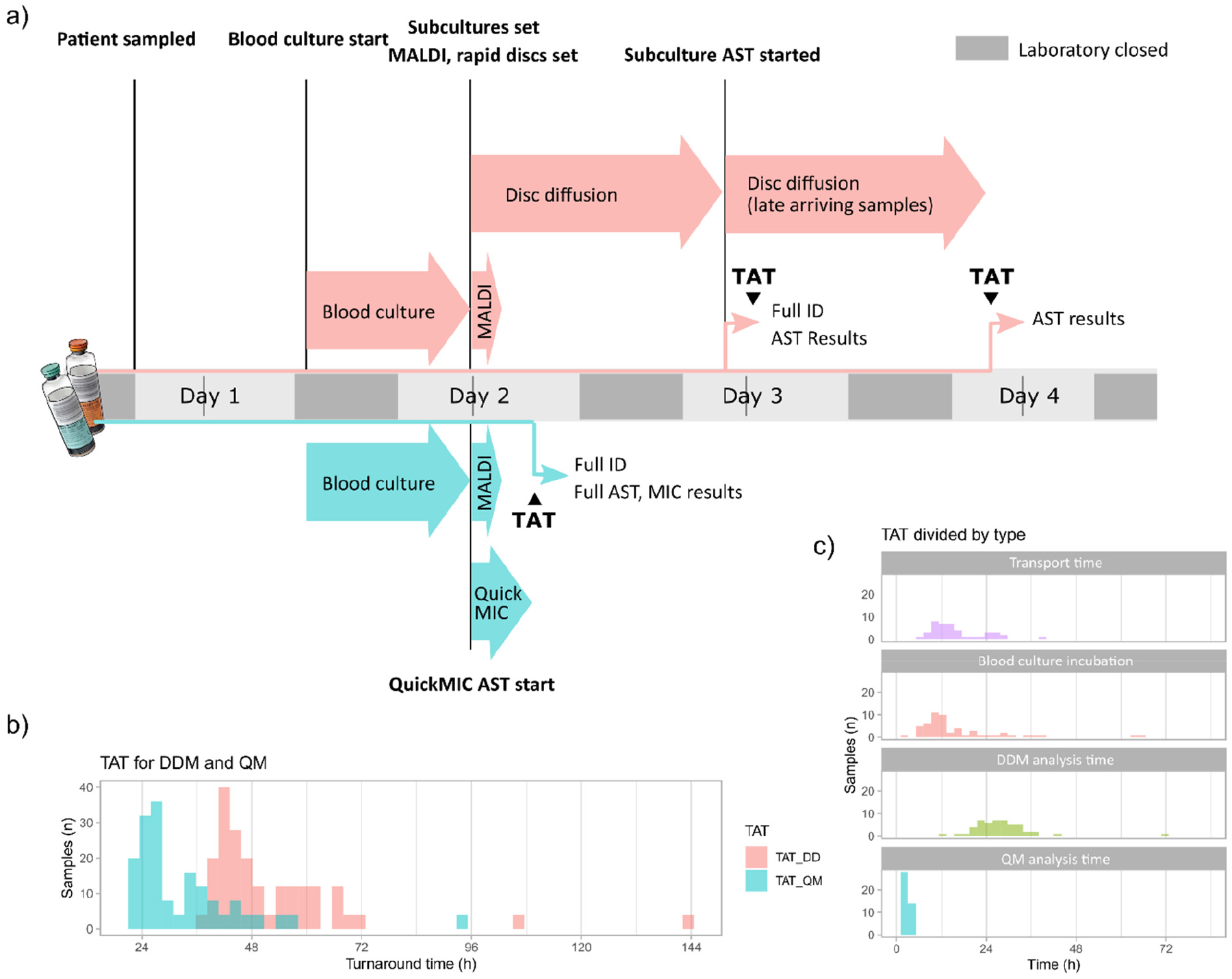
Study layout and turnaround time (TAT). a) The study layout, with tentative timeline for one sample. After patient sampling and sample transport, the blood culture incubation was started. After positivity and during working hours, confirmed Gram-negative samples were run on QuickMIC as well as by the routine laboratory standard process (Maldi-TOF, rapid DDM followed by DDM from isolate). b) The TAT and CA between QuickMIC and DDM were compared. c) AST analysis time, blood culture analysis time and transport times during the study.

## 4 Discussion

This study demonstrates that the QuickMIC system very rapidly can provide AST data for up to 12 antibiotics against at least 10 different species of Gram-negative bacteria. The average time to result was in line with our previous study^15^ with a mean time of ∼3 hours until result, which is significantly shorter than traditional methods such as BMD and DDM. Even though the analysis time of the QuickMIC system is nearly matched by the recently introduced EUCAST rapid disc diffusion (with read-out at 4, 6 and 8 hours), the data resolution is higher and more akin to standard disc diffusion, which we believe is a better comparative method. This study indicates that applying the QuickMIC assay can shorten turnaround times by at least 40% compared to disc diffusion. While the manual nature of rapid disc diffusion read-out constrains read-out times to when a trained microbiologist is available, the automated QuickMIC system could report at any time during the day. Furthermore, the QuickMIC system generates high-resolution quantitative MIC values, whereas rapid disc diffusion provides qualitative susceptibility categories for a limited subset of species and antibiotics.

One limitation of the QuickMIC method is the relatively narrow range of the linear concentration interval provided by the generated antibiotic concentration gradients, which is a trade-off by design to increase the resolution. As a result, very sensitive or very resistant strains will be reported as below or above the quantification limit and not get an on-scale MIC value. We argue that the increased resolution, leading to an increased repeatability as compared with BMD and other log2-based endpoint assays, may allow more accurate results for challenging strains with susceptibilities around the breakpoints. The downside is in situations where clinical breakpoints differ by a large degree between species, which may lead to off-scale breakpoints for a specific panel of antibiotics. Since this method is supposed to be used before species identity is available, it is important to ensure that the range of drug concentrations is wide enough to provide AST results for likely pathogens. In cases where diverging breakpoints exist, one solution can be to include the same antibiotic spanning different concentration ranges in the QuickMIC cassette. In the currently tested Gram-negative panel we do not believe this to be necessary, as all breakpoints for the here tested species are on-scale.

The performance of rapid AST methods is typically determined by either biological factors (such as state of the bacterial culture, innate bacterial growth rate, kinetics of the antibiotic effect) or technical factors (sensitivity and resolution of the system used to measure inhibition). The biological factors can to a certain extent be “optimized”; the culture can be sampled “ready to go” in actively growing state, culture media can be optimized for growth, while the kinetics of antibiotic effects on the other hand cannot be influenced. As for the technical factors, increasing sensitivity and resolution of the system can only improve results up until hitting the limits on time imposed by the biological factors, as measuring with higher sensitivity or much faster will not help if the phenotypic response to the antibiotic is delayed; and slower-growing bacteria will need relatively longer measurement times, regardless of the technology used to measure growth. At the same time, a robust system must be able to handle a wide variety of starting material. It is not practical to mandate from the user to sample the culture at a very specific growth state or concentration. We believe the here presented system in principle represents the fastest possible (i.e. that we have reached the biologically imposed limits) for Gram-negative phenotypic AST under the conditions found in routine clinical microbiology; namely to support a wide range of starting inoculates (from 10^5^ to 10^8^ cfu/mL), growth states (from exponential to stationary phase after multiple hours incubation beyond positive signal), growth rates (from *E. coli* to *P. aeruginosa*) and several routinely used antibiotics (from early-acting bactericidal such as colistin to delayed effect bacteriostatic such as amikacin).

It is also important to note that the main goal when validating new, rapid AST methods is typically to accurately predict the antibiotic susceptibility as measured by a reference method. It is widely known that early antibiotic effects observed in time-kill assays poorly reflect the late effects. Therefore, a measured rapid “MIC_2h_” is likely to differ from the “MIC_18-24h_”, also when using the same method with similar conditions. The challenge of rapid AST is to use the observed MIC_2h_ data to infer the likely MIC_18-24h_ result. The QuickMIC method accomplishes this by quantifying the growth rates and morphological features of every individual bacterial microcolony throughout the antibiotic gradient chamber, thus providing a highly information-dense dataset of the antibiotic-bacteria interaction which maximizes the chance of predicting a correct MIC_18-24h_. Even so, early assessment of AST is not possible for certain combinations of antibiotics, bacterial species, and resistance mechanisms. This problem is shared with other rapid AST methods, and the viability of any new method will be dependent on whether the benefits outweigh the risks, such as an increase in false readouts.

It should be noted that significant factors for a long turnaround time are sample transport times in different parts of the workflow^17^. Logistical improvements such as increased laboratory opening hours and improved sample flows are comparatively easy measures with potentially high impact^17^. Furthermore, we believe that new rapid AST methods need to simultaneously improve on speed but also enable usage scenarios that can assist in shortening sample transport and waiting times. Key to this is automation, modularity, size and usability; where a small and automated system potentially could be located nearer to the patient. It is clear from existing data that one of the main challenges in rapid diagnostics is the logistics of sample handling and opening hours of the laboratory. This is evident in this study as well, where 48.7% of the total turnaround time was represented by the “transport” time category. To reduce these times in time-critical diagnostics such as sepsis, hospitals have started to locate blood culturing cabinets on-site, e.g. at clinical chemistry departments who usually have around the clock opening hours, or even in satellite laboratories nearer to the patient^17^. This is a laudable effort, but the main problem remains: the sample once positive still has to be transported to the laboratory for further analysis. We believe that truly rapid diagnostics will require small, modular and automated AST and ID solutions located near the blood culture cabinets, thus allowing on-site staff such as nurses to start analyses around the clock. We further believe that the here presented QuickMIC system could act as the AST component of such a setup. Incidentally, the same properties can also be argued to be beneficial when examining overall system costs for diagnostic systems in healthcare systems in low- and middle-income countries^13,14^. In such settings, automated analysis systems with high per-sample cost but low overall system cost could be an economically sound alternative to high fixed-cost centralized laboratories.

In summary, we conclude that the QuickMIC system can provide very rapid antibiotic susceptibility results of up to twelve antibiotics per test, directly from positive blood culture samples. The method is accurate, precise and has properties which could allow shorter sample transport chains. By providing rapid AST results, the assay could allow an earlier switch to appropriate targeted therapy, thereby enhancing the chances of survival in critically ill patients and reducing unnecessary use of broad-spectrum antibiotics. However, more studies are needed to assess the clinical benefits of rapid AST in general, and of the QuickMIC system in particular.

## 6 Author Contributions

Author contributions (CRedIT categories) follow: Conceptualization, C.M., CJ, T.T., and J.K.; Data curation, C.M., HÖ.; Formal analysis, C.M.; Funding acquisition, T.T. and J.K.; Investigation, J.T., J.F., C.J. and C.M.; Methodology, C.J., and C.M.; Project administration, C.J and C.M,; Resources, T.T. and J.K.; Software, H.Ö.; Supervision, T.T. and J.K.; Validation, C.M. and C.J.; Visualization, C.M.; Writing – original draft, C.M.; Writing – review & editing, C.M., C.J., T.T., and J.K.

## 8 Acknowledgments

The authors wish to thank Gunnar Kahlmeter and Erika Matuschek at the EUCAST Development Laboratory for fruitful discussions, test strains, and BMD susceptibility data; as well as Sofia Persson, Ehsan Ghaderi and other staff at the Uppsala University Hospital, Department of Clinical Microbiology for support, blood samples and analysis data.

## 9 Data Availability Statement

The datasets analyzed for this study can be provided on request.

